# Functional biological paths altered in Alzheimer’s disease: from genes to bile acids

**DOI:** 10.1101/2020.01.31.929554

**Authors:** Priyanka Gorijala, Kwangsik Nho, Shannon L. Risacher, Rima Kaddurah-Daouk, Andrew J. Saykin, Jingwen Yan

## Abstract

Large-scale genome wide association studies (GWASs) have been performed in search for risk genes for Alzheimer’s disease (AD). Despite the significant progress, replicability of genetic findings and their translation into targetable mechanisms related to the disease pathogenesis remains a challenge. Given that bile acids have been suggested in recent metabolic studies as potential age-related metabolic factors associated with AD, we integrated genomic and metabolomic data together with heterogeneous biological networks and investigated the potential cascade of effect of genetic variations to proteins, bile acids and ultimately AD brain phenotypes. Particularly, we leveraged functional protein interaction networks and metabolic networks and focused on the genes directly interacting with AD-altered bile acids and their functional regulators. We examined the association of all the SNPs located in those candidate genes with AD brain imaging phenotypes, and identified multiple AD risk SNPs whose downstream genes and bile acids were also found to be altered in AD. These AD related markers span from genetics to metabolomics, forming functional biological paths connecting across multiple-omics layers, and give valuable insights into the underlying mechanism of AD.

## Introduction

Alzheimer’s disease (AD) is a neurodegenerative disease characterized by progressive deterioration of memory and other cognitive functions, followed by loss of ability to perform daily activities. It has been affecting an increasing number of aging populations and has now become a public health crisis due to no validated disease modifying treatment. Genetic factors have been widely acknowledged to play a major role in sporadic late onset AD (LOAD). Large-scale twin studies estimated the heritability of AD to be 60-80% [1]. Substantial effort has been dedicated to genome wide association studies (GWASs) in search of AD risk genes. In addition to e4 allele in *APOE*, many other risk variants have been discovered to be involved in AD etiology and many of them are later replicated in other studies, e.g., *BIN1*, *CLU*, *CRI*, *PICALM*, and *CD33* [2–5]. Despite such significant progress, our knowledge about the functions of these risk variants is still very limited. There is a big gap between genetic factors and disease manifestation. How these risk genetic variation leads to the functional impairment in the transcriptomic, proteomic and metabolic levels, and ultimately phenotypic alterations are largely unknown. Therefore, the translation of GWAS findings into targetable mechanisms related to the disease pathogenesis remains a challenge [6].

Converging evidence suggests a key role of cholesterol metabolism in AD. The plasma cholesterol levels in AD are found to be 10% higher than in healthy controls [7]. *APOE*, the top risk gene of AD, is a major regulator of cholesterol metabolism in both peripheral blood and central nervous system [8]. Many other top AD susceptibility loci identified in large scale GWASs are also involved in the cholesterol metabolism (e.g., *APOC1, BIN1, CLU, PICALM, ABCA7*, and *SORL1*) [1, 9, 10]. Bile acids (BAs), a structurally related group of metabolites derived from cholesterol, have been suggested in recent metabolic studies as potential age-related factors associated with AD [11, 12]. BAs are known to be synthesized in the liver and are further chemically modified by the gut microbiota in the distal small intestine and colon. Recent studies in animal models suggest a role for gut microbiome in motor deficit, neuroinflammation [13, 14], cortical beta-amyloid plaque and AD pathogenesis [15]. In addition, there is increasing evidence for the notion that bile acids and the proteins involved in their synthesis are also present in the brain [16–18]. The enzyme that plays a key role in bile acid synthesis, *CYP27A1*, was found to be expressed in the brain and leads to neurological disorders when mutated [19]. In a recent AD study, Nho et. al. found altered bile acid levels in mild cognitive impairment and AD patients [9]. While all of studies suggest potential connections between genetic and bile acid markers, the functional mechanism through which the genetic factors exert effect on the downstream bile acids and ultimately the brain phenotypes is less explored.

In this paper, we hypothesize the cascade of effect of genetic variation on gene expression levels, bile acids concentration, and ultimately brain imaging phenotypes during the development of AD. Integrating genomic and metabolomic data with heterogeneous biological networks, our analysis is focused on the genes directly interacting with AD-altered bile acids and their functional regulators. We examined the additive and epistasis effects of all the SNPs located in those candidate genes on AD-related brain imaging phenotypes and identified 85 AD risk SNPs, where some genes are differentially expressed in AD compared to cognitively normal controls from gene expression analysis in an independent cohort. Taken together, our results revealed that functional biological paths connecting across multi-omics layers can be potentially disrupted by genetic variation in AD. Compared to traditional studies, our novel approach provide additional values to AD research by enabling the discovery of functionally connected markers, and will help provide novel insight into the underlying disease mechanism.

## Materials and Methods

### Study samples

Structural MRI and GWAS genotype data analyzed in the present report were obtained from the Alzheimer’s disease Neuroimaging Initiative (ADNI) cohort. The initial phase (ADNI-1) was launched in 2003 to test whether serial magnetic resonance imaging (MRI), position emission tomography (PET), other biological markers, and clinical and neuropsychological assessment could be combined to measure the progression of mild cognitive impairment (MCI) and early AD. ADNI-1 was extended to subsequent phases (ADNI-GO, ADNI-2, and ADNI-3) for follow-up for existing participants and additional new enrollments. Inclusion and exclusion criteria, clinical and neuroimaging protocols, and other information about ADNI can be found at www.adni-info.org [10, 20]. Demographic information, raw neuroimaging scan data, GWAS genotype data, and clinical information are available and were downloaded from the ADNI data repository (www.loni.usc.edu/ADNI/). Written informed consent was obtained at the time of enrollment that included permission for analysis and data sharing and consent forms were approved by each participating sites’ Institutional Review Board (IRB).

### MRI imaging processing

Magnetic Resonance Imaging (MRI) T1-weighted brain MRI scans at baseline were acquired using a sagittal 3D MP-RAGE sequence following the ADNI MRI protocol [20, 21]. As detailed in previous studies, FreeSurfer V5.1, a widely employed automated MRI analysis approach, was used to process MRI scans and extract whole brain and ROI (region of interest)-based neuroimaging endophenotypes including volumes and cortical thickness determined by automated segmentation and parcellation [22, 23] The cortical surface was reconstructed to measure thickness at each vertex. The cortical thickness was calculated by taking the Euclidean distance between the grey/white boundary and the grey/cerebrospinal fluid (CSF) boundary at each vertex on the surface [24–26]. In total, 73 cortical thickness measures and 26 volume measures from different brain regions were extracted. 20 of them presented in Table 1 were found to be highly relevant to AD [27] and were used as quantitative traits in this analysis.

**Table 1.**
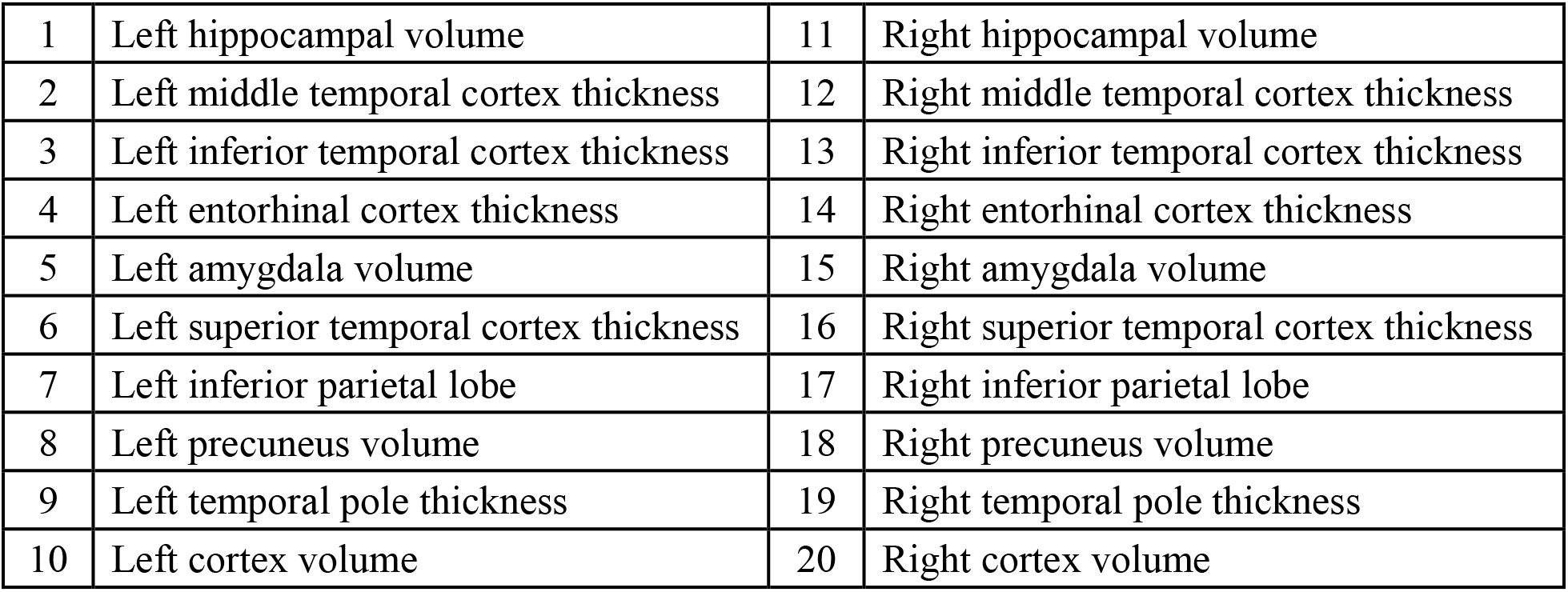
List of 20 AD related imaging phenotypes.

|  |  |  |  |
| --- | --- | --- | --- |
| 1 | Left hippocampal volume | 11 | Right hippocampal volume |
| 2 | Left middle temporal cortex thickness | 12 | Right middle temporal cortex thickness |
| 3 | Left inferior temporal cortex thickness | 13 | Right inferior temporal cortex thickness |
| 4 | Left entorhinal cortex thickness | 14 | Right entorhinal cortex thickness |
| 5 | Left amygdala volume | 15 | Right amygdala volume |
| 6 | Left superior temporal cortex thickness | 16 | Right superior temporal cortex thickness |
| 7 | Left inferior parietal lobe | 17 | Right inferior parietal lobe |
| 8 | Left precuneus volume | 18 | Right precuneus volume |
| 9 | Left temporal pole thickness | 19 | Right temporal pole thickness |
| 10 | Left cortex volume | 20 | Right cortex volume |

### GWAS genotype data preparation

ADNI samples were genotyped using Human 610-Quad, HumanOmni Express, and HumanOmni 2.5M BeadChip. Sample and SNP quality control procedures of GWAS data such as SNP call rate < 95%, Hardy-Weinberg equilibrium test p < 1e-6, and frequency filtering (MAF ≥ 5%) were performed. Only non-Hispanic Caucasian participants were selected by clustering with CEU (Utah residents with Northern and Western European ancestry from the CEPH collection) + TSI (Toscani in Italia) populations using HapMap 3 genotype data and the multidimensional scaling (MDS) analysis after performing standard quality control procedures for genetic markers and subjects. Un-genotyped SNPs were imputed separately using the 1000 Genomes Project-based reference panel as the ADNI cohort used different genotyping platforms.

### Bile acid data preparation

Targeted metabolomics profiling was performed to identify and quantify concentrations of 20 bile acids (BA) from serum samples using Biocrates® Bile Acids Kit as described in detail [9]. In brief, morning serum samples from the baseline visit were collected and aliquoted as described in the ADNI standard operating procedures, with only fasting samples included in this study [28]. BA quantification was performed by liquid chromatography, tandem mass spectrometry. Metabolites with >40% of measurements below the lower limit of detection were excluded. To assess the precision of the measured analytes, a set of blinded analytical replicates (24 pairs in ADNI-1 and 15 triples in ADNI-GO/2) were supplied by ADNI. Unblinded metabolite profiles went through further quality control (QC) checks. The preprocessed dataset included 15 BAs that passed QC criteria. The preprocessed BA values obtained from the QC step were adjusted for the effect of medication use (at baseline) on BA levels [29]. Among that, nine bile acids were found to be significantly associated with AD in a recent study [27], where AD-related brain imaging phenotypes were used as quantitative traits.

### Network integration

We manually curated metabolic networks and protein interaction networks from existing databases (Fig 1A). First, we downloaded the functional protein interactions from the REACTOME database [30], which consists of predicted functional interactions and a large portion of interactions extracted from expert-curated pathways [30]. The interactions were included only if they have 1) direction information (e.g., activation or inhabitation) because we are interested in the upstream genetic regulators of bile acids and 2) confidence scores greater than 0.7 if they are predicted.

**Fig 1.**
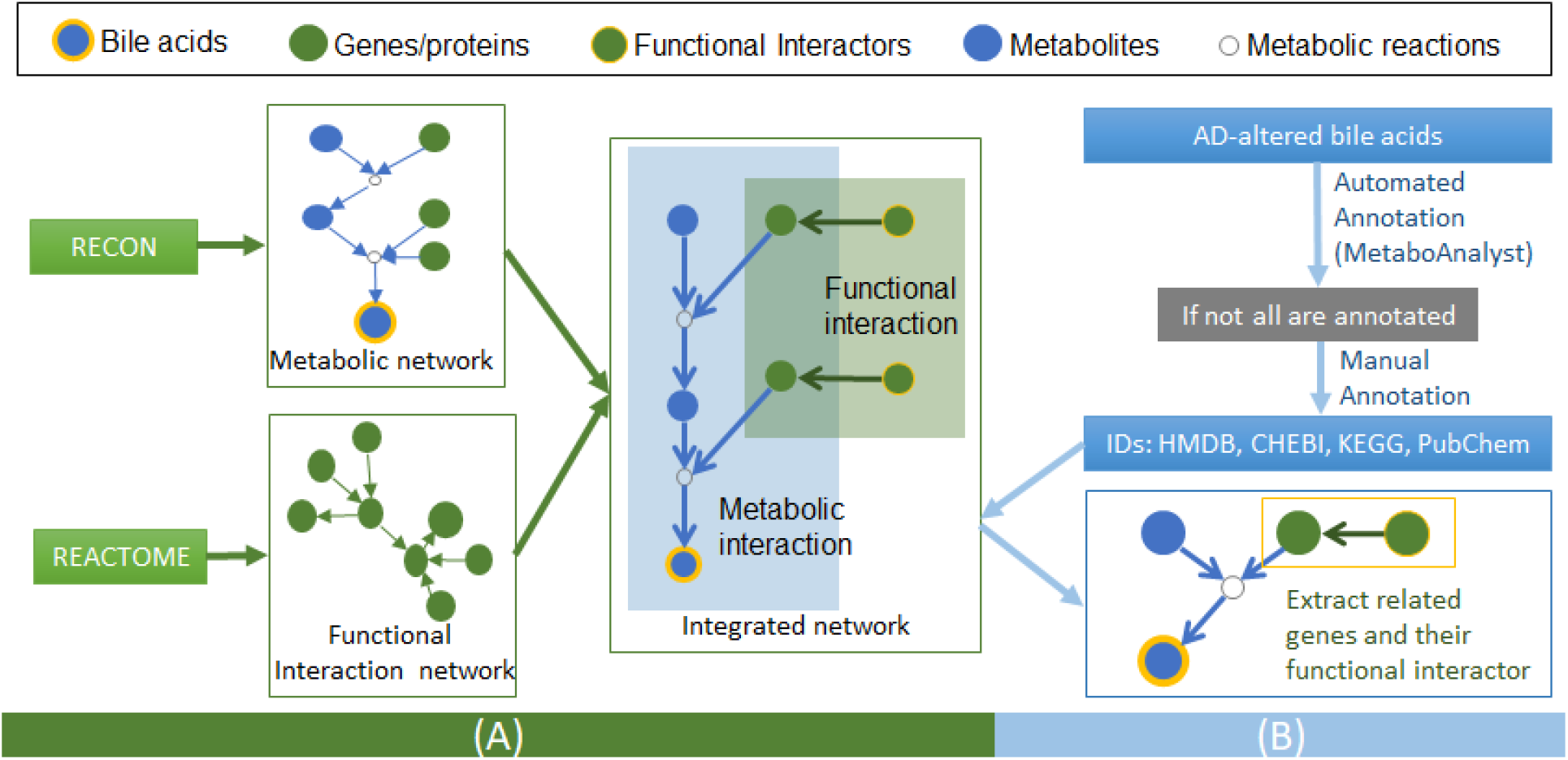
Workflow for network integration (A) and extraction (B) of genes related to AD-altered bile acids.

Second, we downloaded metabolic reactions from RECON [31, 32]. It is now the most comprehensive human metabolic reconstruction available to us, which details all known metabolic reactions occurring in human. All reactions in RECON were curator-validated. We extracted the gene-reaction and metabolite-reaction relationships from their raw data and combined into a metabolic network involving genes, metabolites and reactions. For each reaction, there is one input metabolite, one output metabolite and one or more coding genes for enzymes, receptors or transporters. Finally, we integrated the functional interaction network with the metabolic network based on Entrez ID to generate an integrative network characterizing multiple layers of regulation relationships.

### Extraction of genes related to bile acids

A list of candidate genes functionally related to nine AD-altered bile acids were extracted based on the integrated network (Fig 1B). To map the bile acids onto the network, we first applied an automated approach, MetaboAnalyst (www.metaboanalyst.ca/), to annotate the bile acids and search for their unique identifiers, such as HMDB ID, PubChem Compound ID, ChEBI ID, or KEGG ID. The bile acids that could not be automatically matched to existing database entries were manually annotated. We went through the literatures for their synonyms and searched PubChem, KEGG and CHEBI websites for detailed structure information to confirm and cross-reference the IDs. For bile acids with at least one identifier, we mapped them to the integrated network and extracted a list of candidate genes including enzymes, receptors and transporters directly related to them. The candidate gene set was further expanded by including their upstream genetic interactors (i.e., functional regulators in the REACTOME protein interaction network).

### Targeted genetic association analysis

We extracted a set of SNPs from all candidate genes within ±50 kb of each gene boundary and tested their association with 20 AD-related brain imaging phenotypes using a linear regression model in Plink [33]. Age, gender, education, MRI field strength, intracranial volume (ICV) and presence/absence of *APOE* ε4 allele were used as covariates. Given that both SNPs and brain imaging phenotypes are highly correlated, we estimated the number of linkage disequilibrium (LD) blocks as *N*_*L*_ and the number of independent phenotypes as *N*_*P*_. The number of independent association tests is calculated as *N*_*L*_ * *N*_*P*_. On top of this, we applied the Bonferroni Correction procedure for multiple comparison adjustment.

### Targeted genetic interaction analysis

We further examined the epistasis effect of all SNPs on 20 AD-related brain imaging phenotypes using epistasis model in Plink [33]. Age, gender, education, MRI field strength, intracranial volume (ICV) and presence/absence of *APOE* ε4 allele were used as covariates. All epistasis results were Bonferroni corrected for the estimated number of independent interaction tests (i.e.,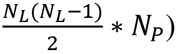.

### Transcriptomic association analysis

For SNPs significantly associated with brain phenotypes from temporal cortex regions (in either GWAS association or epistasis analysis), we identified their corresponding genes and compared their expression levels between cognitively normal controls and AD using RNA-Seq data generated from the temporal cortex tissue in the Mayo Clinic cohort [34]. If one gene is not directly interacting with the AD-altered bile acids, but through a linker gene, the expression pattern of these linker genes was also examined. The differential expression analysis was performed using R package, EdgeR [35], where age, sex, batch, RIN and *APOE* ε4 status were used as covariates. EdgeR is one of the most robust packages for differential expression analysis, which uses a generalized linear model (GLM) approach based upon negative binomial distribution. Normalization of the RNA-Seq data was performed using Trimmed mean of M values (TMM method). Prior to analysis, genes were filtered with counts per million as very low counts provide little evidence for differential expression.

## Results

### Study sample after QC

After QC procedures, 1,416 ADNI participants with GWAS genotype and structural MRI data at baseline were used, including 325 healthy controls (HC), 87 significant memory concerns (SMC), 261 early mild cognitive impairment (EMCI), 465 late MCIs (LMCI), and 278 AD patients. Detailed Demographic information for the study samples is presented in Table 2.

**Table 2.** Demographic and clinical characteristics of ADNI participants.

|  | HC | SMC | EMCI | LMCI | AD |
| --- | --- | --- | --- | --- | --- |
| Number | 325 | 87 | 261 | 465 | 278 |
| Gender(M/F) | 165/160 | 36/51 | 143/118 | 287/178 | 156/122 |
| Mean age | 73.37(7.19) | 72.08(7.24) | 73.36(7.20) | 73.37(7.19) | 73.32(7.20) |
| Mean education | 15.96(2.84) | 16.37(2.71) | 15.95(2.85) | 15.95(2.84) | 15.94(2.85) |
| *APOE* (ε4-/ ε4+) | 234/90 | 58/29 | 150/110 | 211/254 | 94/184 |
| Shown in the parentheses are the standard deviations. |  |  |  |  |  |

### Selection of candidate genes

Out of nine bile acids associated with AD, we were able to map four onto the integrated network, including chenodeoxycholic acid (CDCA), glycolithocholic acid (GLCA), glycodeoxycholic acid (GDCA), and taurolithocholic acid (TLCA). For these 4 bile acids, we extracted genes directly interacting with them, as enzymes, receptors or transporters, and also indirect genetic interactors, which connect to the bile acids through a linker gene. In total, we identified 23 candidate genes related to four AD-altered bile acids, including 14 direct interactors and 9 indirect ones (Fig. 2). Based on the reference genome, 4,964 SNPs, belonging to 157 LD blocks, were identified within ±50kb from the boundary of these candidate genes.

**Fig 2.**
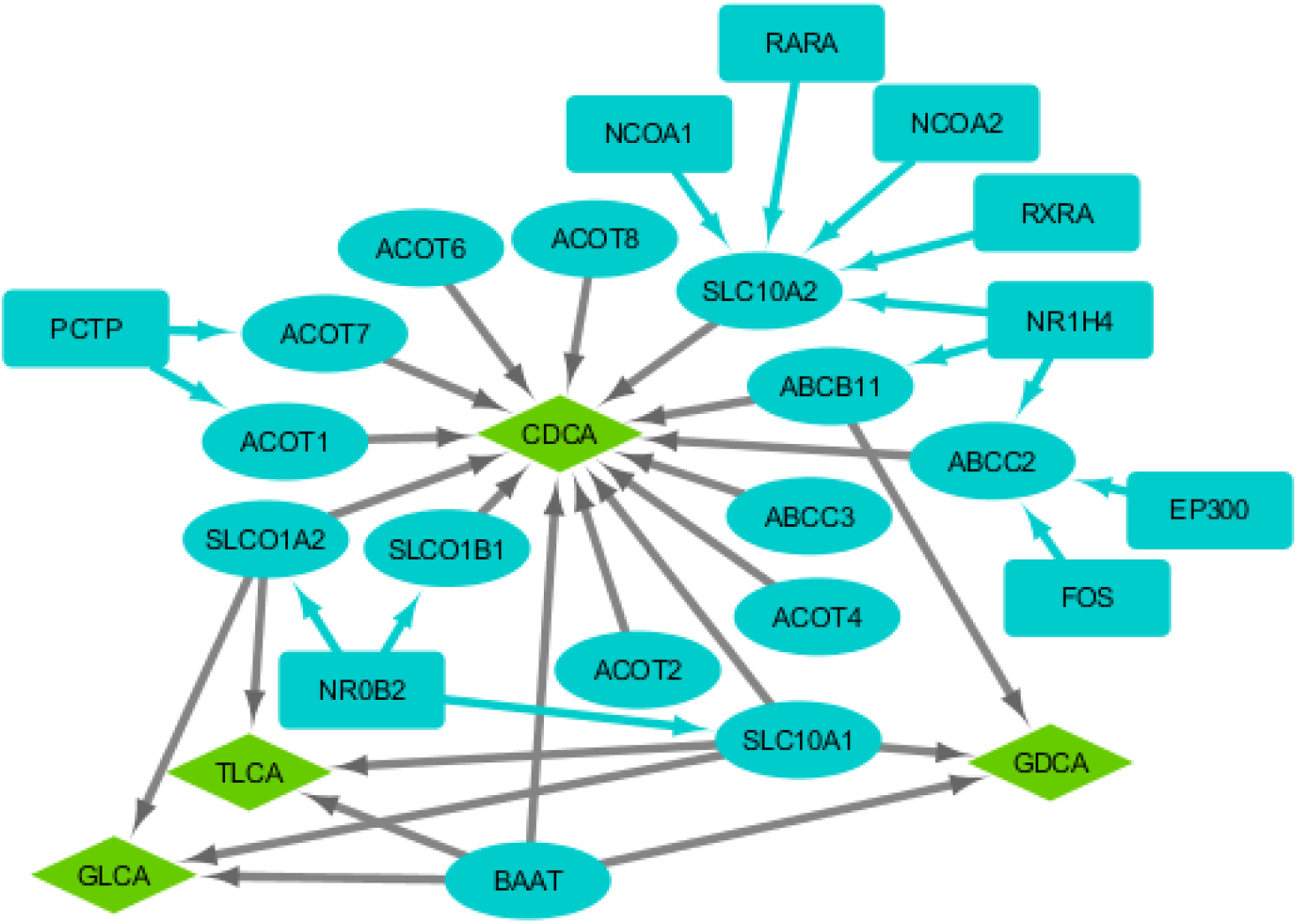
Candidate genes included in our analysis and their interaction with bile acids. Green: bile acids; Blue (rectangular): genes indirectly interacting with bile acids; Blue (ellipse): genes directly interacting with bile acids; Gray links: metabolic interactions; Blue links: functional protein interactions.

### Targeted genetic association analysis

Linear regression models were used to examine the association between each pair of SNPs and brain imaging phenotypes. The number of LD blocks is estimated as *N*_*L*_ = 157 and 20 brain imaging phenotypes is found to fall into 3 blocks (*N*_P_ = 3) (Fig. 3). After correcting for the estimated number of independent test, our association analysis identified 29 SNPs as significantly associated with left entorhinal cortex thickness and right precuneus volume (Fig. 4). More specifically, these SNPs include rs116159950, rs117327561, rs141853576 in *NR1H4* belonging to distinct LD blocks, and 26 SNPs from the same LD block of *ACOT7*. To better understand the function of these two genes, we further examined their connection with AD risk genes [3] in the functional interaction network using Cytoscape Reactome FI plugin (Fig. 5). While *ACOT7* is located in the downstream of AD risk genes, *NR1H4* is predicted to interact with *APOE* through the linker gene *ESR1*. According to our integrated network (Fig. 2), *NR1H4* regulates *SLC10A2, ABCB11,* and *ABCC2*, which mediate the synthesis of CDCA and/or GDCA, two bile acids associated to AD [27].

**Fig 3.**
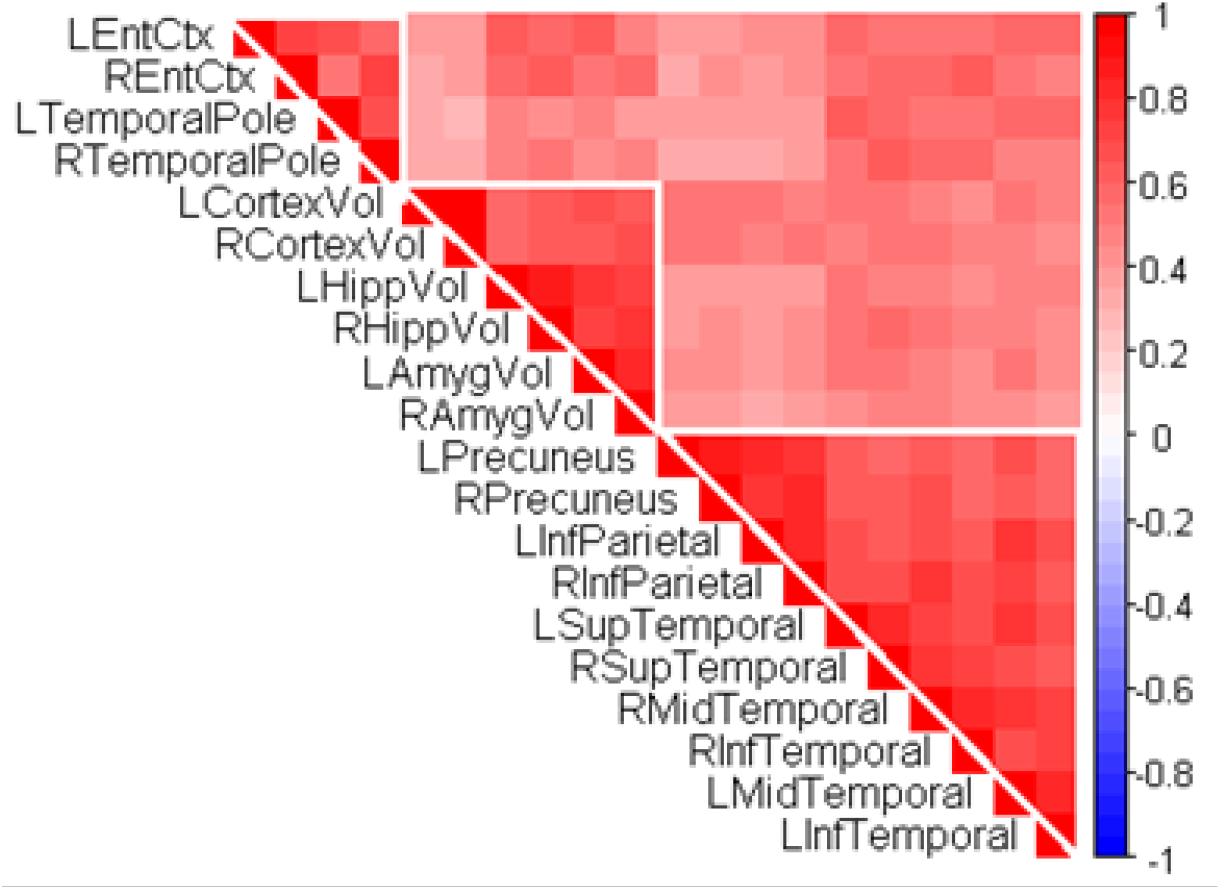
Association results of 20 AD-related brain imaging phenotypes.

**Fig 4.**
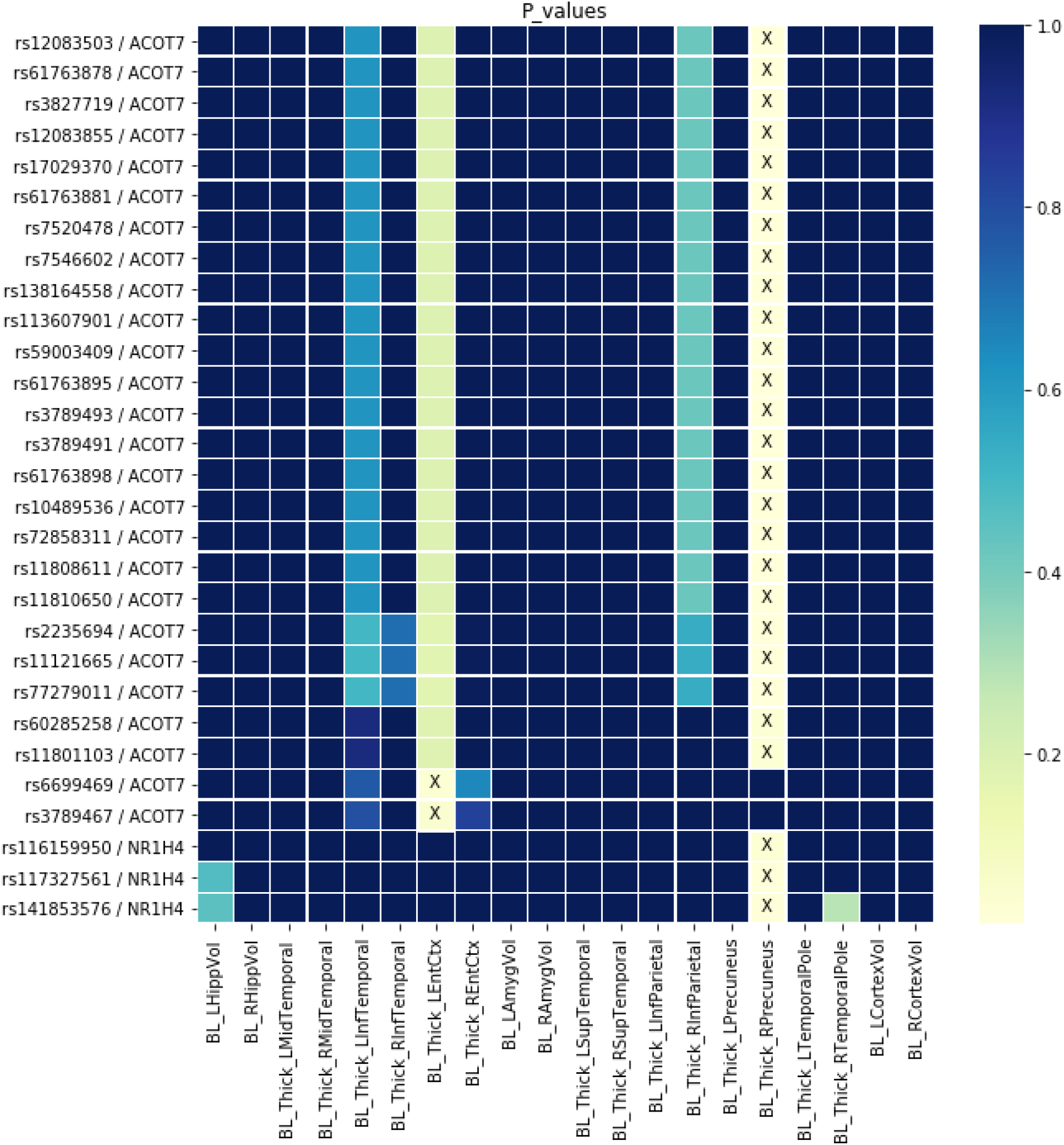
Significant GWAS associations identified between SNPs and AD-related brain phenotypes. Crossed portions indicate associations with corrected p-value<0.05.

**Fig 5.**
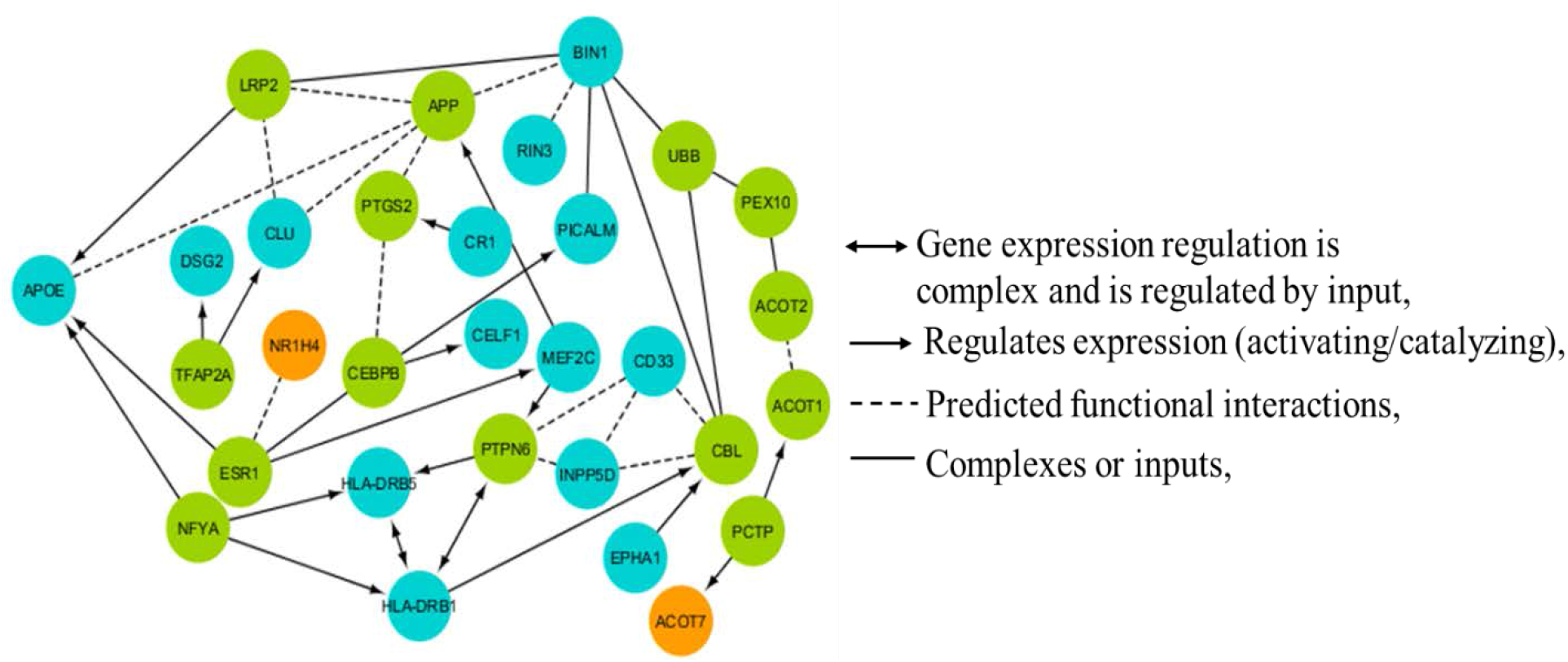
Functional interaction network between *ACOT7*,*NR1H4* and AD-risk genes. Green: linker genes; Blue: AD risk genes; Orange: genes identified as significant from targeted GWAS association analysis.

### Epistasis analysis

Genetic interaction of all SNPs was examined against each of 20 brain imaging phenotypes. We estimated the number of independent interaction tests as 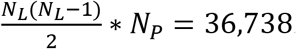. After Bonferroni correction, 5,337 SNP pairs were found to be significant (See Supplementary File S1). These are 85 unique SNPs in nine genes, including *ACOT7, EP300, ACOT8, NCOA1, RARA, SLC10A1, NR1H4, SLCO1A2, and ABCC2*. Significant interactions were observed in the right inferior parietal, right inferior temporal, right mid temporal, right precuneus, right superior temporal, right temporal pole regions. Shown in Fig. 6 is the distribution of significant interactions across brain phenotypes. The majority of the significant genetic interactions was found in the temporal cortex region. The interaction between rs76524143 (*NCOA1*) and rs8036602 (*EP300*), showed the lowest p value (3.59E-08) after multiple testing correction.

**Fig 6.**
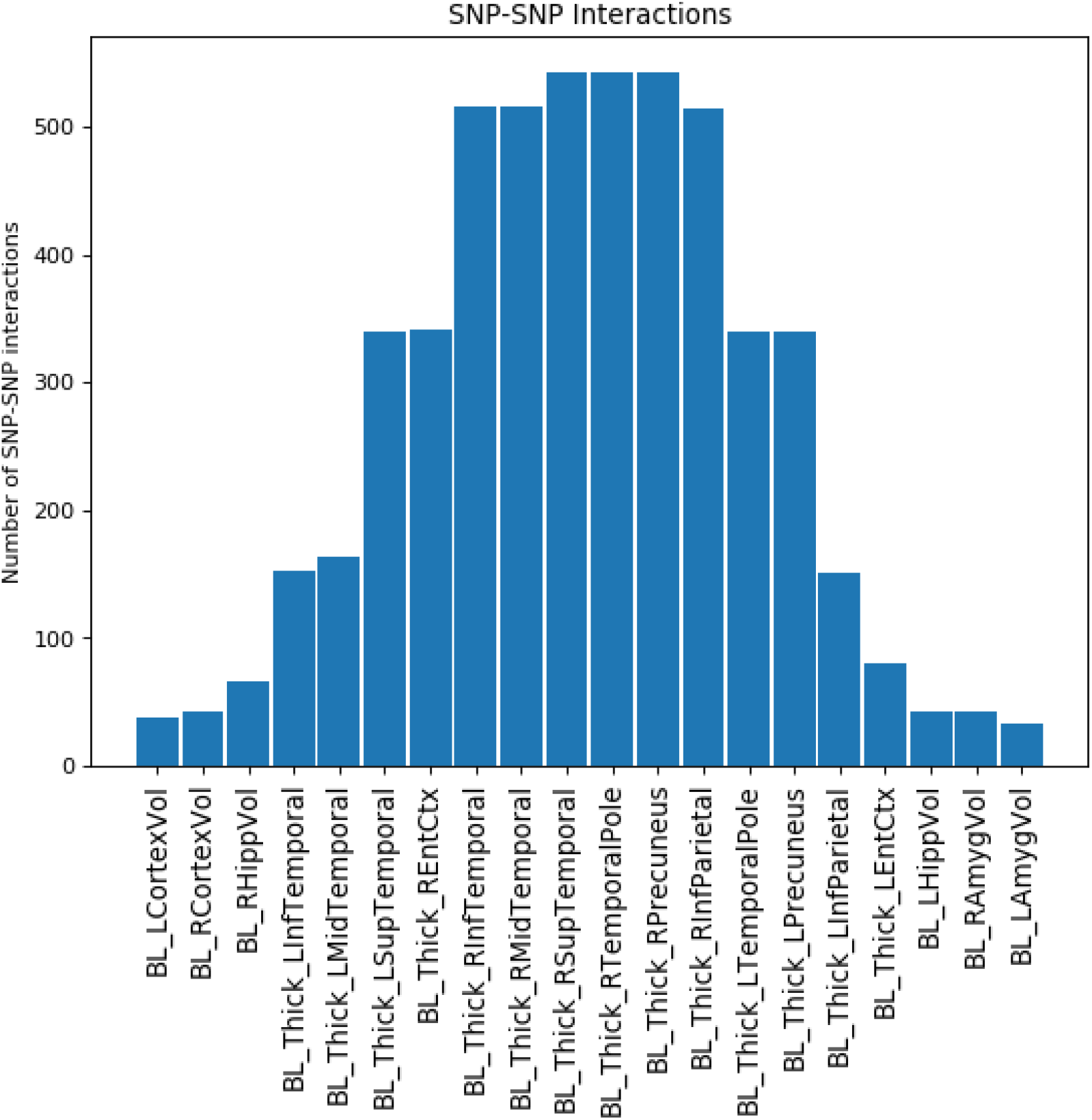
Distribution of significant SNP-SNP interactions in all 20 phenotypes.

To better understand the function of the genes hosting these SNPs, we further examined the connection between these genes and AD risk genes [3] in the functional interaction network using Cytoscape Reactome FI plugin (Fig. 7). All 9 genes are closely connected with AD risk genes. In particular, *EP300* shows as a hub in the functional interaction network, regulating 14 AD risk genes directly or indirectly through linker genes. According to our integrated network (Fig 2), *EP300* functionally regulates *ABCC2* gene, which mediates the synthesis of bile acid CDCA.

**Fig 7.**
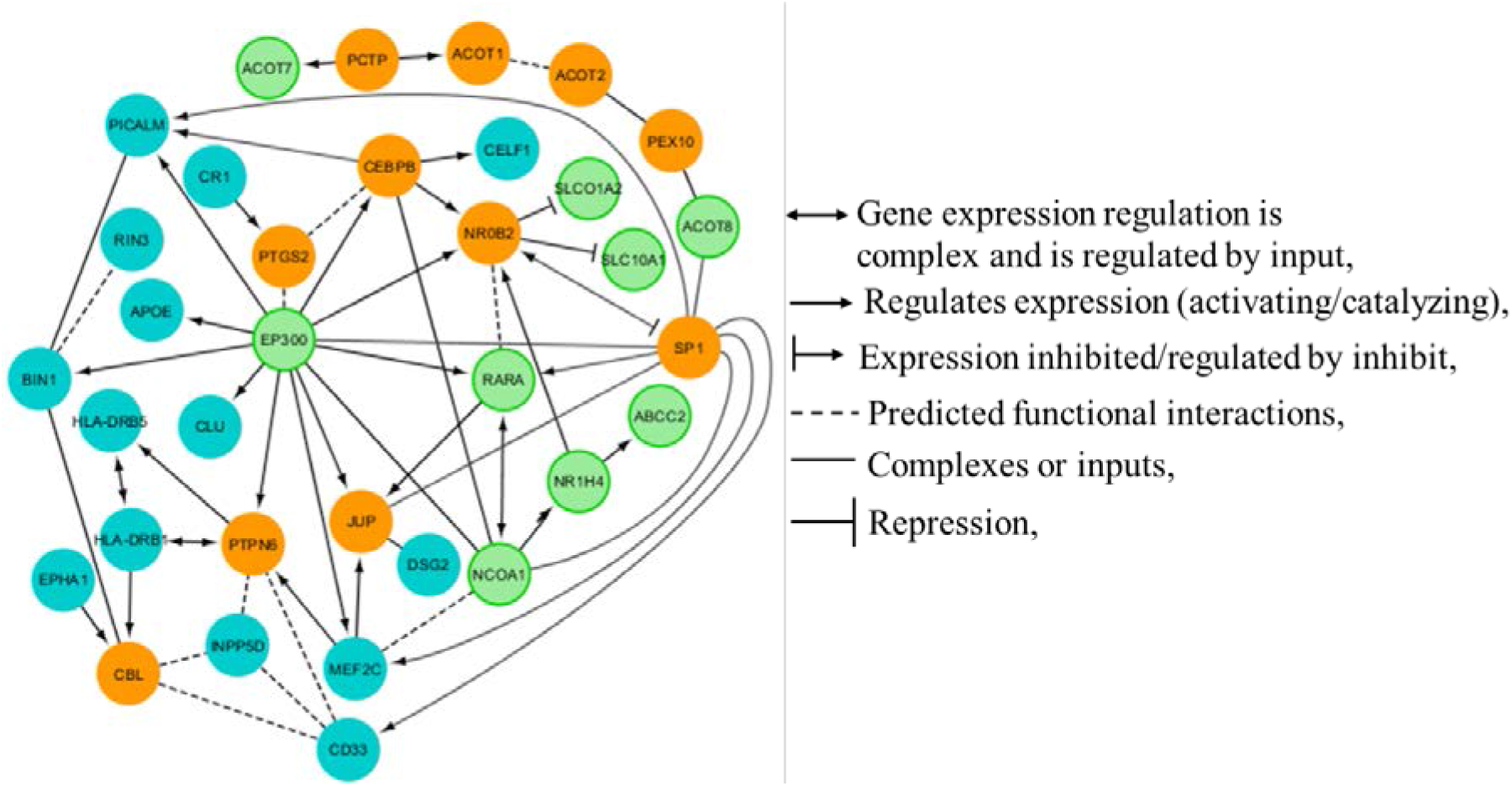
Functional interaction network between AD-risk genes and significant genes identified in epistasis analysis. Blue: AD risk genes, Green: significant genes from epistasis results, Orange: linker Genes.

### Differential expression of significant genes in brain

For all SNPs showing significant associations with imaging phenotypes related to the temporal cortex regions in SNP-based analysis or epistasis analysis, we identified their corresponding genes and compared their expression patterns between cognitively normal controls and AD using RNA-Seq data generated from the temporal cortex region [34]. If one gene does not interact with AD-altered bile acids directly, but through a linker gene, expression levels of the linker gene were also examined. Out of 10 genes, we identified 4 as significantly differentially expressed in the temporal cortex region after multiple testing correction. A summary of differentially expressed genes (DEGs) was presented in Table 3.

**Table 3.**
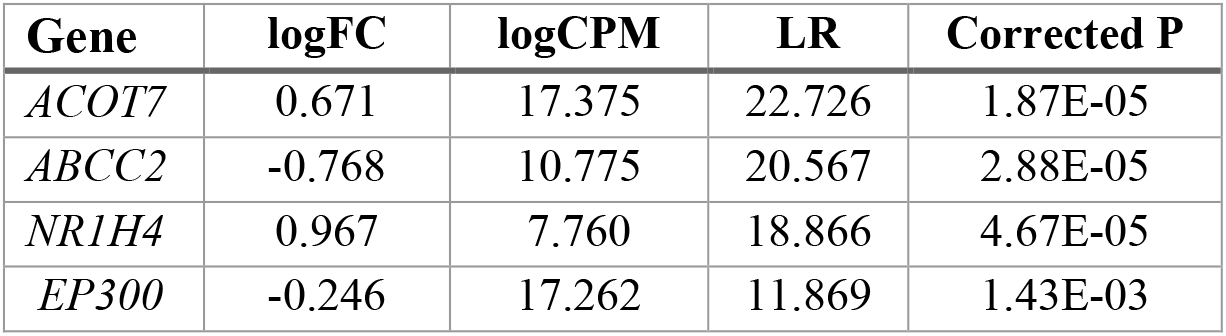
Summary of differentially expressed genes.

## Discussion

In this study we explored the potential cascade effect of genetic variations on the downstream genes, bile acids and AD quantitative traits. Results from this study revealed a set of heterogeneous AD markers, forming functional biological pathways that cut through multi-omics layers from genome to transcriptome and phenome. For example, as shown in the epistasis results, the upstream regulators, nuclear receptor subfamily 1 group H member 4 (*NR1H4)* and histone acetyltransferase p300 (*EP300)* both harbor SNPs significantly associated with the thickness of temporal cortex regions. In temporal cortex, transcription factor *NR1H4* and its direct binding targets ATP binding cassette subfamily C member 2 (*ABCC2),* which directly mediate the level of CDCA, were both found differentially expressed in AD patients (Fig. 8).

**Fig 8.**
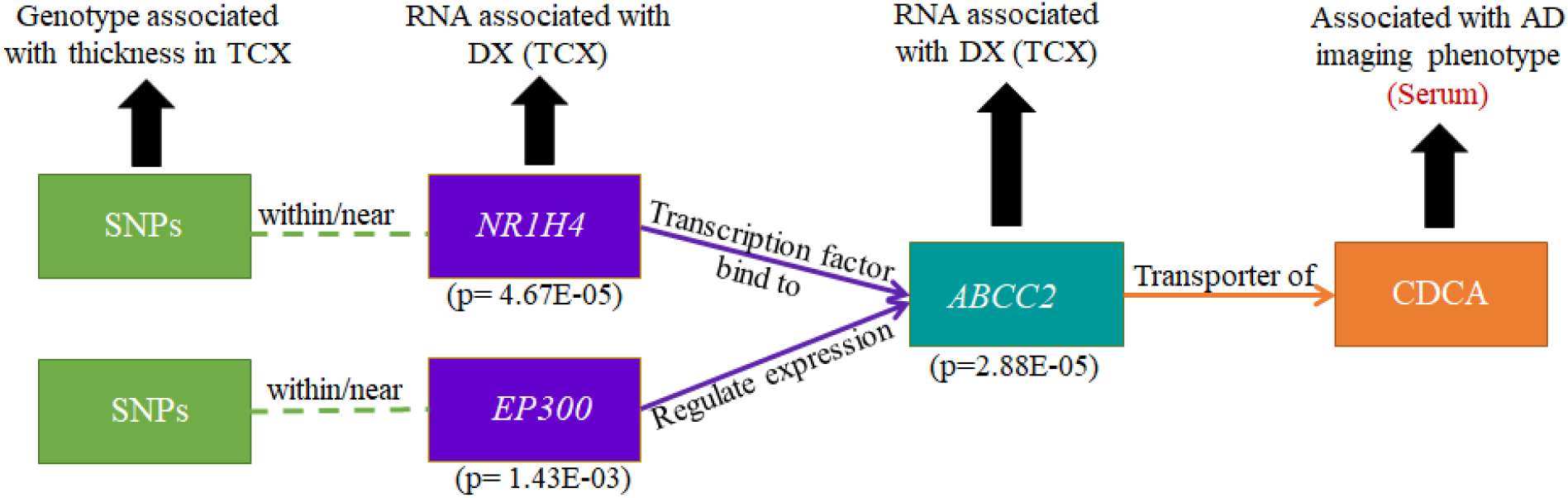
A subset of omics network summarizing our results. DX: diagnosis. TCX: temporal cortex.

The results of this study were further supported by the reported role of *NR1H4/FXR* (Farnesoid X receptors) in cholesterol homeostasis. *NR1H4/FXR* is commonly expressed in the liver, intestine and kidneys. It was reported that *FXR* agonists, such as bile acids and their derivatives, were found to be useful in metabolic syndrome therapies. Later, Huang et al identified functional *FXR* in brain neurons. Their result provided evidence of the presence of *FXR* mRNA and protein in the freshly isolated human brain cortex and hippocampal regions, and also potential transcriptional activity of *FXR* [36]. *NR1H4* has been suggested as a potential therapeutic target to slow down AD progression using potent *FXR* agonists. Studies show that inhibition of *FXR* signaling help reduce the severity of neurological disorders allied with over accumulation of bile acids [37, 38].

*EP300* gene encodes the enzyme histone acetyltransferase p300 or E1A-associated protein p300, also known as *EP300* or *P300*. This enzyme functions as histone acetyltransferase that regulates transcription of genes via chromatin remodeling. Findings from multiple studies have suggested the potential of *P3*0*0* to act as a biomarker for dementia assessment and monitoring AD. Meta-analysis of *P300* amplitude and latency reveals useful information about the early stages of AD [39, 40]. The brain regions typically affected in AD, such as frontal cortex parietal cortex and hippocampus, generate *P300* wave. It was also strongly associated with cognitive variables, which are impaired in AD as well [41]. In addition, *P300* is one of the top candidate master regulators effecting the transcriptional signature of AD progression. The activity of *P300* acetyltrasferase was found to be upregulated in AD brain tissue. Acetylation homeostasis dysregulation, one of the possible reasons for neurotoxicity, is mediated by *P300* through hyperacetylation of its substrates p53 and histone3 proteins, which are significantly increased in AD brain tissue [42].

*ABCC2*/*MRP2*, regulated by both *NR1H4* and *EP300*, are responsible for the transportation of bile acids. Findings from existing studies suggest that ABC transporters are involved in amyloid β accumulation in aging brain and have a potential to serve as futuristic targets in AD therapy [43, 44]. A recent microarray study on AD and brain aging suggested that the regulator of *ABCC2*, *EP300*, as a Ca^2+^ dependent incipient AD regulating gene [45] and altered Ca^2+^ signaling was suspected to have a role in AD [46].

*ACOT7*, also known as brain acyl-CoA hydrolase, was previously shown to be enriched in the brain [47, 48]. It is an important regulator in neurometabolism and protects neurons against the damaging effects of fatty acid excess [49]. Loss of *ACOT7* expression results in multiple defects in the regulation of fatty acid metabolism in the brain. It is suggested as vital regulatory point in neuronal fatty acid homeostasis. An improved understanding of *ACOT7* is beneficial to protect against neurotoxicity owing to the fact that neurological disorders arise from metabolic dysfunction (i.e., lipid metabolism dysregulation). Furthermore, *ACOT7* was found to be significantly upregulated in the late stage of AD [50].

## Conclusion

In this study, we performed an integrative analysis of genotype, bile acids and brain imaging phenotype data by leveraging the functional interactions between proteins and bile acids as priori. We identified several AD risk genetic variation, whose downstream genes and bile acids are found to be altered in AD or associated with AD. Taken together, this suggests several trans-omic biological paths disrupted during the development of AD, due to the cascade effect of genetic variation. To the best of our knowledge, this is the first study that analyzed multi-omics data from different layers and generated functional biological paths as potential system biomarkers for AD.

## Acknowledgments

Funding for ADMC (Alzheimer’s Disease Metabolomics Consortium, led by Dr R.K.-D. at Duke University) was provided by the National Institute on Aging grant R01AG046171, a component of the Accelerated Medicines Partnership for AD (AMP-AD) Target Discovery and Preclinical Validation Project (https://www.nia.nih.gov/research/dn/amp-ad-target-discovery-and-preclinical-validation-project) and the National Institute on Aging grant RF1 AG0151550, a component of the M2OVE-AD Consortium (Molecular Mechanisms of the Vascular Etiology of AD – Consortium https://www.nia.nih.gov/news/decoding-molecular-ties-between-vascular-disease-and-alzheimers).

Data collection and sharing for this project was funded by the Alzheimer’s Disease Neuroimaging Initiative (A.D.N.I.) (National Institutes of Health Grant U01 AG024904) and DOD A.D.N.I. (Department of Defense award number W81XWH-12–2-0012). A.D.N.I. is funded by the National Institute on Aging, the National Institute of Biomedical Imaging and Bioengineering, and through generous contributions from the following: AbbVie, Alzheimer’s Association; Alzheimer’s Drug Discovery Foundation; Araclon Biotech; BioClinica, Inc.; Biogen; Bristol-Myers Squibb Company; CereSpir, Inc.; Eisai Inc.; Elan Pharmaceuticals, Inc.; Eli Lilly and Company; EuroImmun; F. Hoffmann-La Roche Ltd and its affiliated company Genentech, Inc.; Fujirebio; GE Healthcare; IXICO Ltd.; Janssen Alzheimer Immunotherapy Research & Development, LLC.; Johnson & Johnson Pharmaceutical Research & Development LLC.; Lumosity; Lundbeck; Merck & Co., Inc.; Meso Scale Diagnostics, LLC.; NeuroRx Research; Neurotrack Technologies; Novartis Pharmaceuticals Corporation; Pfizer Inc.; Piramal Imaging; Servier; Takeda Pharmaceutical Company; and Transition Therapeutics. The Canadian Institutes of Health Research is providing funds to support A.D.N.I. clinical sites in Canada. Private sector contributions are facilitated by the Foundation for the National Institutes of Health (www.fnih.org). The grantee organization is the Northern California Institute for Research and Education, and the study is coordinated by the Alzheimer’s Disease Cooperative Study at the University of California, San Diego. A.D.N.I. data are disseminated by the Laboratory for Neuro Imaging at the University of Southern California.

This research was also supported by NIH grants R21 AG066135, R01 EB022574, R01 AG019771, P30 AG010133, NSF grant 1755836, NLM R01 LM012535, NIA R03 AG054936, Indiana University Collaborative Research Grant (IUCRG) and Enhanced Mentoring Program with Opportunities for Ways to Excel in Research (EMPOWER). This project was also funded, in part, with support from the Indiana Clinical and Translational Sciences Institute funded, in part by Grant Number UL1TR001108 from the National Institutes of Health, National Center for Advancing Translational Sciences, Clinical and Translational Sciences Award.

## Supporting information

**S 1 File. Significant SNP-SNP interactions of each brain phenotype.**

